# Direct cortical stimulation induces short-term plasticity of neural oscillations in humans

**DOI:** 10.1101/2023.11.15.567302

**Authors:** Saachi Munot, Naryeong Kim, Yuhao Huang, Corey J. Keller

## Abstract

Patterned brain stimulation is commonly employed as a tool for eliciting plasticity in brain circuits and treating neuropsychiatric disorders. Although widely used in clinical settings, there remains a limited understanding of how stimulation-induced plasticity influences neural oscillations and their interplay with the underlying baseline functional architecture. To address this question, we applied 15 minutes of 10Hz focal electrical simulation, a pattern identical to ‘excitatory’ repetitive transcranial magnetic stimulation (rTMS), to 14 medically-intractable epilepsy patients undergoing intracranial electroencephalographic (iEEG). We quantified the spectral features of the cortico-cortical evoked potential (CCEPs) in these patients before and after stimulation. We hypothesized that for a given region the temporal and spectral components of the CCEP predicted the location and degree of stimulation-induced plasticity. Across patients, low frequency power (alpha and beta) showed the broadest change, while the magnitude of change was stronger in high frequencies (beta and gamma). Next we demonstrated that regions with stronger baseline evoked spectral responses were more likely to undergo plasticity after stimulation. These findings were specific to a given frequency in a specific temporal window. Post-stimulation power changes were driven by the interaction between direction of change in baseline power and temporal window of change. Finally, regions exhibiting early increases and late decreases in evoked baseline power exhibited power changes after stimulation and were independent of stimulation location. Together, these findings that time-frequency baseline features predict post-stimulation plasticity effects demonstrate properties akin to Hebbian learning in humans and extend this theory to the temporal and spectral window of interest. These findings can help improve our understanding of human brain plasticity and lead to more effective brain stimulation techniques.

**Significance Statement:** Brain stimulation is increasingly used to treat neuropsychiatric disorders by inducing changes in neural activity at specific brain regions. Despite their effectiveness, how these changes occur, specifically in the spectral domain, is unknown. To better understand how brain oscillations change after patterned stimulation, we performed focused stimulation in epilepsy patients and measured intracranial brain recordings. We found strong and predictable changes in brain oscillations (plasticity) after patterned stimulation. Specifically, low frequencies showing widespread effects and high frequencies exhibiting a greater magnitude of change. These changes were directly related to the temporal and spectral structure of brain responses prior to stimulation. Our study reveals that baseline brain activity patterns can predict how stimulation will induce plasticity in the spectral domain. These findings can help improve our understanding of human brain plasticity and lead to more effective brain stimulation techniques.

**Highlights:**

1. We applied 15 minutes of repetitive 10Hz focal electrical stimulation and assessed the evoked brain-wide spectral changes with intracranial EEG.
2. 10Hz stimulation induced short-term plasticity in low frequency alpha evoked power broadly across regions and time windows and high frequency (beta, gamma) power specifically in early evoked time windows (10-50ms).
3. Across patients, frequency bands, and time windows, brain regions with stronger baseline evoked power were more likely to undergo greater spectral changes after 10Hz stimulation.
4. Post-stimulation spectral changes were specific; that is, for a given frequency band in a specific time window, baseline evoked power predicted post-stimulation change in the same frequency band and time window.
5. Post-stimulation spectral change was driven by an interaction between direction of change and temporal window of baseline power; that is, regions exhibiting baseline evoked early (10-100ms) increases and late (100-200ms) decreases in power correlated with observed post-stimulation spectral changes.
6. These results were independent of stimulation location.

## Introduction

Neuropsychiatric disorders are some of the most common and debilitating disorders in the world (Whiteford et al., 2015). Brain stimulation treatments including repetitive transcranial magnetic stimulation (TMS) are clinically effective alternatives to medications (Garnaat et al., 2018 & Croarkin et al., 2021). Despite their efficacy, the neural mechanisms underlying stimulation-induced clinical changes remain unclear. Studying why specific brain regions and neural activity change after stimulation can help elucidate the neural mechanism underlying clinical effects and harness principles of neuroplasticity to optimize and deliver more effective treatment.

Despite this need for increased understanding, novel methods are needed to elucidate the neural effects of repetitive stimulation in humans. With respect to stimulation, TMS elicits sensory effects that can confound the interpretation of brain responses (Conde et al., 2019). On the recording side, while non-invasive brain measures such as functional MRI (fMRI) and electroencephalography (EEG) can provide some insight, these tools lack high temporal (fMRI) or spatial (EEG) resolution, respectively. In this investigation, we overcome these limitations by applying focal electrical stimulation and recording brain responses using intracranial EEG (iEEG). Focal electrical stimulation does not typically elicit perceptual changes and thus bypasses the sensory confounds of TMS, while iEEG provides high spatial and temporal resolution of neural activity (Ross et al., 2022; Lucas et al., 2023). In this study we probe evoked oscillatory changes after repetitive stimulation using corticocortical evoked potentials (CCEPs). CCEPs have been used to predict the onset of ictal events (David et al., 2013), examine the functional brain architecture (Keller et al., 2011, 2014b; David et al., 2013; Entz et al., 2014), and recently quantify changes after repetitive stimulation (Keller et al., 2018 & Huang et al., 2019). This work demonstrated that regions that were anatomically close and functionally connected predicted post-stimulation changes, as probed with CCEPs (Keller et al., 2018; Huang 2019). Importantly, this study did not evaluate how repetitive stimulation modulates time-frequency oscillatory activity that can be extracted from iEEG with high spatiotemporal resolution. This precise time-frequency information would segregate oscillatory effects at specific frequencies of interest, a key physiological dimension which may be foundational to the brain’s functioning (Buzsáki & Draguhn, 2004). Thus, still missing is an understanding of the relationship between evoked oscillatory events prior to stimulation and where, when, and in which frequency bands changes occur after repetitive stimulation. As such, we applied 10 minutes of 10Hz focal repetitive stimulation to multiple brain regions and measured pre/post stimulation changes with CCEPs (Keller et al., 2014, Keller et al., 2018 & Keller et al., 2011). We hypothesized that regions that are functionally connected will undergo neural plasticity after stimulation, and that this concept will extend to the time-frequency domain. That is, the change in functional connectivity will have a temporal and frequency-based specificity. First, after repetitive stimulation we observed group-level changes in evoked oscillations, where across frequency bands regions with stronger baseline power were more likely to undergo post-stimulation changes. Low frequency (alpha) power demonstrated broad post-stimulation effects, while high frequency (beta, gamma) power demonstrated the strongest effects, especially in earlier time windows. As hypothesized, we also observed that baseline spectral features were more likely to correlate with post-stimulation effects in a spatially and temporally specific manner. Together, these results offer insight into the neural mechanisms underlying human neuroplasticity and provide future direction for novel neuromodulatory interventions to enhance neuroplasticity.

## Materials and Methods

### Subjects

Fourteen subjects with medically-intractable epilepsy underwent surgical implantation of intracranial electrodes for seizure localization. Subject characteristics are detailed elsewhere (Keller, Honey, Entz, et al., 2014). Subjects were enrolled at two hospitals: North Shore University Hospital (Manhasset, New York, USA) and National Institute of Clinical Neurosciences (Budapest, Hungary). All subjects provided informed consent as monitored by the local Institutional Review Board and in accordance with the ethical standards of the Declaration of Helsinki. The decision to implant, the electrode targets, and the duration of implantation were made entirely on clinical grounds without reference to this investigation. Subjects were informed that participation in this study would not alter their clinical treatment and that they could withdraw at any time without jeopardizing their clinical care.

### Electrode registration

Our electrode registration method has been described in detail previously (Dykstra et al., 2012 & Groppe et al., 2017). Briefly, in order to localize each electrode anatomically, subdural electrodes were identified on the post-implantation CT with BioImagesuite (Dykstra et al., 2012), and were coregistered first with the post-implantation structural MRI and subsequently with the pre-implantation MRI to account for possible brain shift caused by electrode implantation and surgery. Following automated coregistration, electrodes were snapped to the closest point on the reconstructed pial surface (Dykstra et al., 2012) of the pre-implantation MRI in MATLAB (Dykstra et al., 2012). Intraoperative photographs were previously used to corroborate this registration method based on the identification of major anatomical features. Automated cortical parcellations were used to localize electrodes to anatomical regions (Groppe et al., 2017).

### Electrophysiological recordings

This electrophysiological dataset has been described previously (Keller et al., 2018 & Huang et al., 2019). Here, we provide a brief overview of the data and outline new statistical analyses performed for this work. Invasive electrocorticographic (iEEG) recordings from implanted intracranial subdural grids, strips and/or depth electrodes were sampled at 512 or 2048Hz depending on clinical parameters at the participating hospital (U.S.A. and Hungary, respectively). Data preprocessing and analysis was performed using the FieldTrip toolbox (Oostenveld et al., 2011). Single pulses of direct electrical stimulation induced stereotyped stimulation artifacts that were ∼10ms in duration. To remove pulse artifact, we applied a 4^th^ order bandpass filter (100-150Hz, Butterworth, two-pass) and replaced this time period with stationary iEEG time series that represented a similar amplitude and spectral profile as the background signal, as described previously (Keller et al., 2018, Huang et al., 2019, Crowther et al., 2019). Line noise (60Hz and 50Hz for recordings in the U.S.A and Hungary, respectively) was then removed using a notch filter. Following artifact rejection and line noise removal, we applied a bipolar montage to depth electrodes and a common average reference montage to grid/strip electrodes (Stolk et al., 2018).

### Repetitive stimulation paradigm

We applied focal 10Hz electrical stimulation in a temporal pattern timed to match that of rTMS treatment for depression, as previously described (Keller et al., 2018 & Huang et al., 2019). Each subject received 15 minutes of 10Hz direct electrical stimulation in bipolar fashion (Fig 1A; biphasic pulses at 100 us/phase, 5s trains, 10s rest, 15s duty cycle, total 3000 pulses). Stimulation sites included frontal, temporal, and parietal regions as described in our previous work.

**Fig 1:**
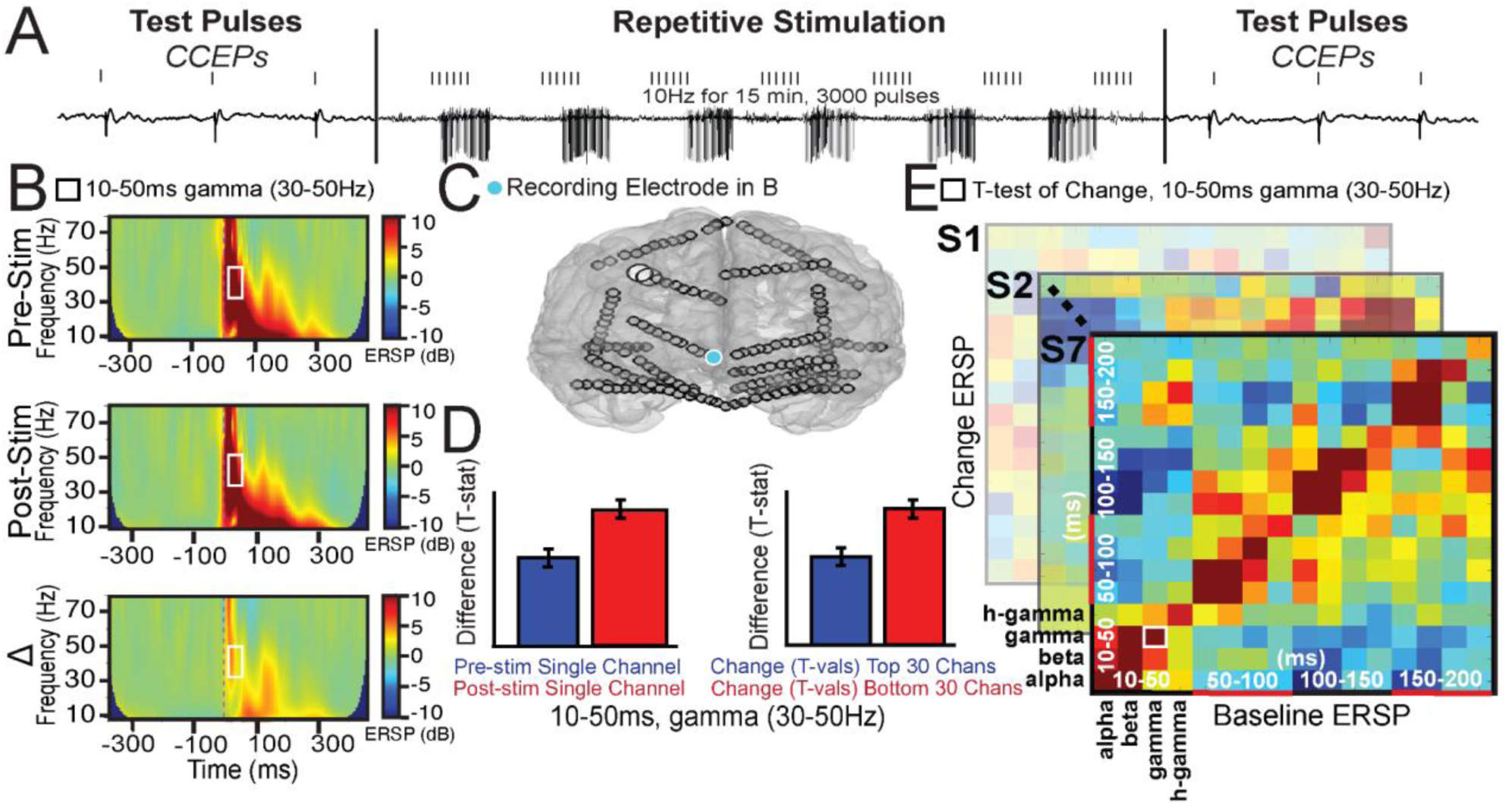
Schematic and workflow. A) Stimulation paradigm. 10Hz direct electrical stimulation, patterned to mimic a single treatment session of rTMS, was applied to induce neural changes. Changes were assessed by comparing CCEPs before and after 10Hz stimulation. B) *Top and Middle:* Single pulse event-related spectral perturbation (ERSP) map before and after 10Hz stimulation, generated from one channel in one subject (S7). The square denotes one time-frequency feature (10-50ms low gamma power). *Bottom:* Change in ERSP before and after stimulation. C) *Top:* Localization of stimulation electrodes (white) and recording electrodes (blue, from ERSP plots in B). D) *Left:* Single channel statistical comparison of pre- vs post-stimulation responses with a single time-frequency feature. The blue bar represents pre-stimulation trials and the red bar represents post-stimulation trials. *Right:* Single subject statistical comparison of pre-stimulation and change (pre vs post) for a single time-frequency feature across channels. E) Heatmaps for each subject. Here, each pixel represents the t-value from the second bar graph in D on the right; that is, comparing baseline to pre/post change for each time-frequency feature.

### Pre/post stimulation CCEPs

To examine causal changes in brain excitability at baseline and after stimulation, we performed cortico-cortical evoked potential (CCEP) mapping (Keller, Honey, Mégevand, et al., 2014, Keller et al., 2017, Keller et al., 2018, Huang et al., 2019). Prior to and immediately following 15 minutes of 10Hz focal electrical stimulation, to assess excitability changes we applied 200 pulses of bipolar electrical stimulation (biphasic pulses at 100 us/phase) with a 1s inter-stimulation interval (ISI). Reasons for inter-stimulus interval (ISI) and jitter timing considerations have been described previously (Keller et al., 2018). This CCEP paradigm in of itself has not been shown to induce neural changes (Keller et al., 2018). Briefly, to evaluate CCEPs in each channel, data were epoched (-1000 to 1000ms) and standardized using Z-scores against the pre-CCEP baseline period (-150ms to -50ms), as detailed previously (Keller et al., 2018). As such, here we use the term *short-term plasticity* to refer to the spectral changes in the CCEP that occur on the order of minutes after 10Hz stimulation.

### Spectral Feature Quantification

To extract oscillatory features from the CCEP before and after 10Hz stimulation, in each channel a wavelet spectral decomposition was performed (Fig 1B) (Delorme & Makeig, 2004). We used the newtimef function for this wavelet analysis (EEGLAB, San Diego, California), with the following parameters: padratio 8, cycles [1.5 0.7] (increasing linearly with frequency), and output frequencies 8-120Hz. Lower frequencies <8 Hz were not computed due to insufficient data points within 250ms of the stimulation pulse that were not contaminated by the stimulation pulse artifact. To ensure results were not specific to the wavelet decomposition approach, we also calculated the spectrogram, and results did not qualitatively differ (see supplementary Fig S1, S2, and S3).

Next, time-frequency features were extracted from the CCEP. Feature extraction included power in each frequency band >8Hz (alpha 8-12Hz, beta 12-25Hz, gamma 25-50Hz, high gamma 50-100Hz) during specific time windows of the CCEP (10-50ms, 50-100ms, 150-200ms). In an effort to capture differences between the N1 (<50ms) and N2 (>50ms) components of the CCEP (Keller et al., 2011, Keller et al., 2014), as well as to extract features in equivalent time windows that can be directly compared, we used the following time bins: 10-50ms, 50-100ms, 100-150ms, and 150-200ms. As described above, data within 10 ms of the stimulation pulse were excluded to avoid stimulation artifacts. To compute power for each time-frequency feature, the absolute value of the processed data was log transformed (20*log10(abs(tfdata))) and baseline corrected. To determine changes in these time-frequency features after 10Hz stimulation, we compared pre- and post-stimulation evoked time-frequency features using a two-sample t-test (ttest2, MATLAB; Fig 1D) for each temporal (i.e. 10-50ms) and spectral (i.e. high gamma) window. The resultant t-value represents the degree of change for each evoked time-frequency after 10Hz stimulation. This strength of post-stimulation change was then compared to the evoked time-frequency features before 10Hz stimulation.

### Relating baseline to post-stimulation oscillatory activity

To test the hypothesis that baseline evoked oscillatory activity predicts changes elicited after 10Hz stimulation, we compared baseline (pre-10Hz) evoked power in the CCEP to the t-value denoting pre/post 10Hz change in spectral-temporal features of the CCEP (as described above in Spectral Feature Quantification Methods section). To do so, for each participant, and time-frequency feature we first removed outliers (+/- 3 SD). To compare baseline power to pre/post change in power, we split baseline power by selecting the top and bottom quartiles to ensure there were enough samples for statistical comparison. For each time-frequency feature of the CCEP, we next compared the pre/post change metric in channels with high and low baseline power, and extracted a second t-value comparing baseline to pre/post change. For each time-frequency feature, the average baseline-to-pre/post change t-value was computed (Fig 1D) and represented as one pixel in the heatmap in (Fig 1E). To normalize for shifts in patient specific t-value distributions, a weighted average analysis was also performed (see supplemental Fig S4). To quantify significant effects for each time-frequency feature, we generated a null distribution. To do so, instead of comparing sets of channels with low vs high pre-stimulation power, we computed the t-statistic from two sets of randomly-selected channels. This process was repeated 200 times for each participant,resulting in 2800 data points (N=14). Finally, from this distribution, we defined the 97.5^th^ and 2.5^th^ percentile as statistically significant (asterisks in Fig 2 and 3).

**Fig 2:**
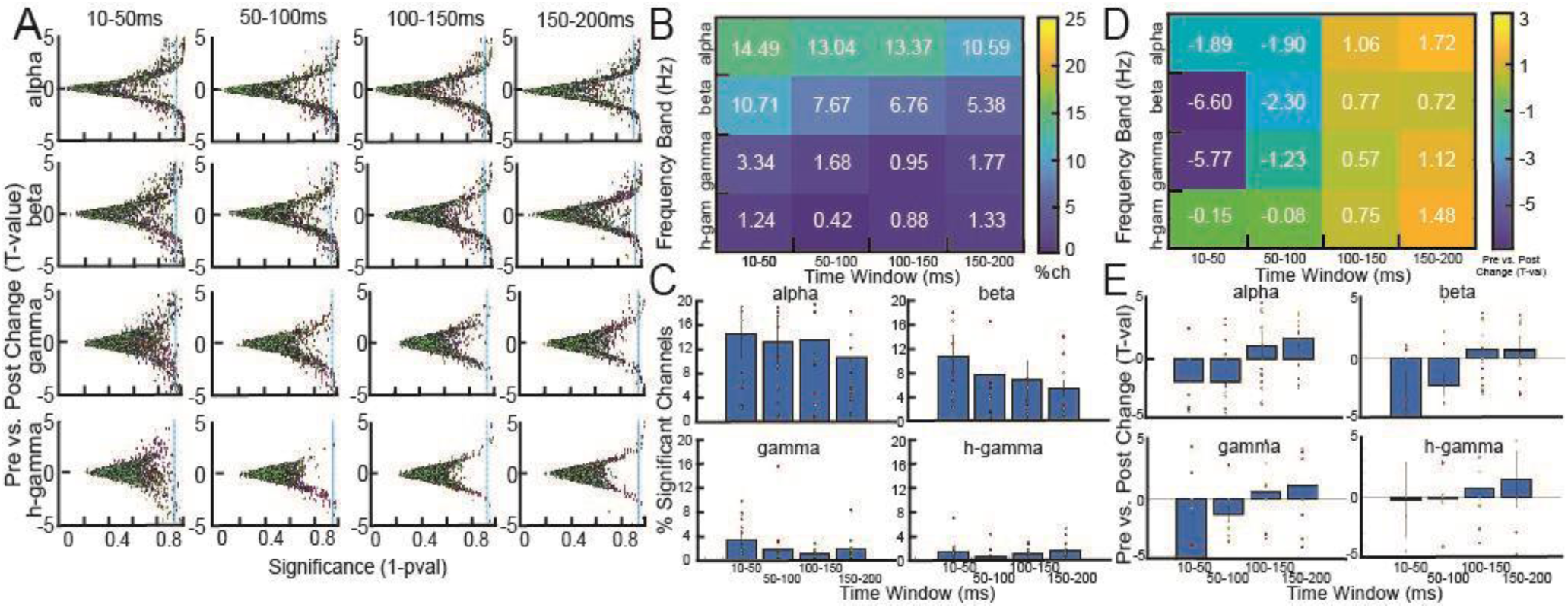
Relationship between spectral features, sensitivity to change and magnitude of change in neural oscillatory activity after repetitive stimulation. A) Distribution of channels for all patients per frequency band and time window relating degree and significance of change. Those on the right of the blue line show significant change. B, C) Heatmap and bar plots showing sensitivity to change, characterized by the group-level percentage of significant channels (p<0.05) for each frequency band and time window. D, E) Heatmap and bar plots showing group-level net magnitude of change across all significant channels.

**Fig 3:**
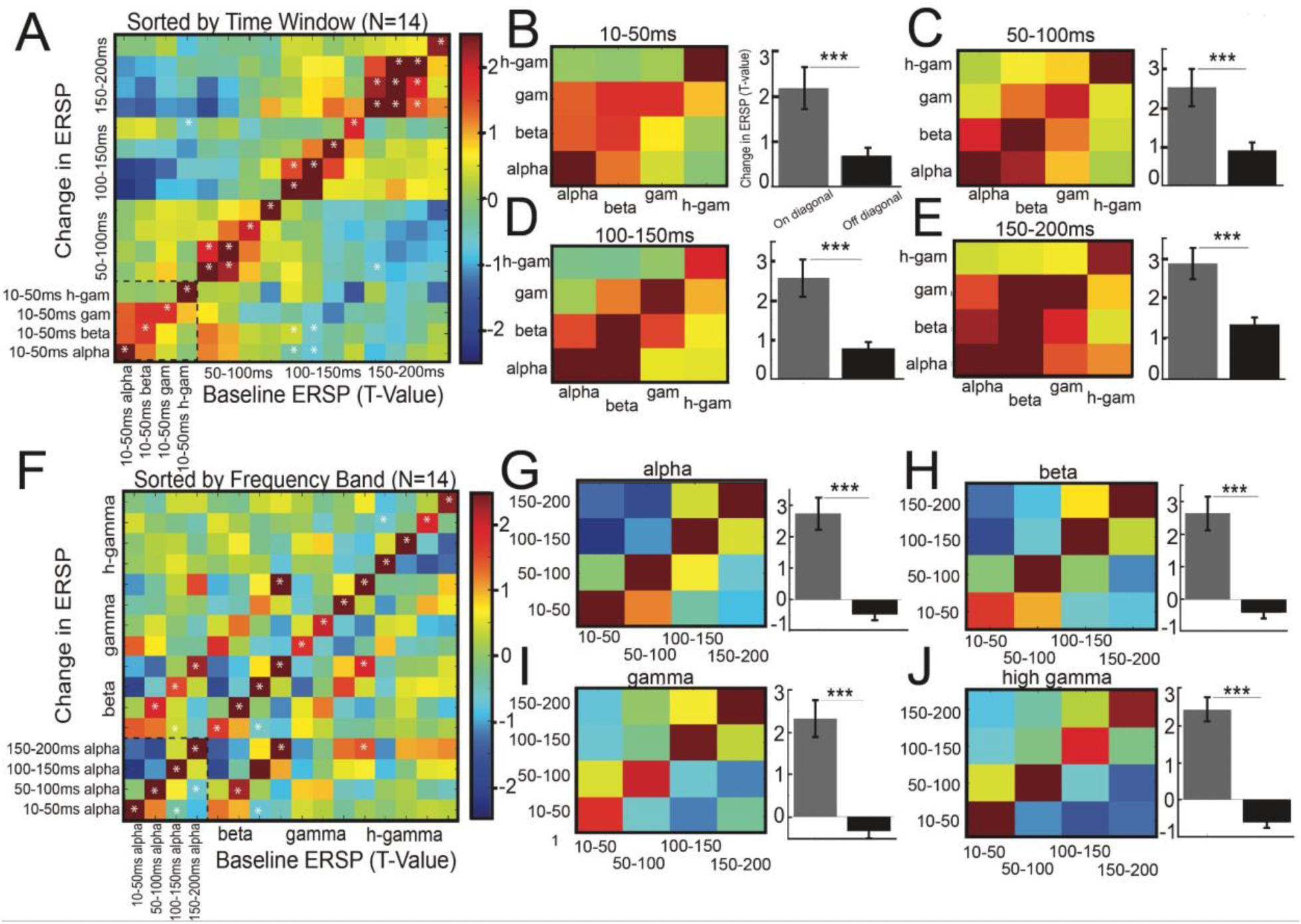
Baseline evoked spectral features predict post-stimulation change after repetitive stimulation. A) Group quantification relating baseline and pre/post change in event-related spectral perturbation (ERSP). Axes are sorted by (A) time window and (F) frequency. Note the strong effect between baseline time-frequency features and the magnitude of change of those features after 10Hz stimulation, as quantified by the diagonal. *p<0.05 from bootstrapped null distribution (see Methods). B-E) Same data from A but visualized within each time window. Bar graphs represent comparison of on-diagonal vs off-diagonal; that is, how baseline time-frequency features predict post-stimulation change in the same time-frequency feature. F) Same as A but sorted by frequency band and latency. G-J) Same as B-E but sorted by frequency band. For bar graphs, *p<.05, **p<.01, ***p<.001 (Student’s T-test)

### Statistical analysis of group-level data

First, to compare the neural effects of pre- and post- 10Hz stimulation, spectral-temporal features of the CCEP were grouped by frequency, time window, and baseline direction of change (e.g., if there was an observed increase or decrease in baseline power for each time-frequency feature, for example gamma 10-50ms after the pulse). To examine the effect of direction of pre-stimulation evoked power on pre/post spectral changes we performed the above analysis stratified by regions exhibiting positive and negative baseline evoked power (Fig 3). For example within the 10-50ms gamma bin, channels exhibiting pre-stimulation evoked gamma power suppression were analyzed separately from those exhibiting pre-stimulation evoked gamma power increases. Furthermore, to examine those channels with strong baseline power, we further stratified by the top and bottom quartile of baseline evoked power. In summary, this analysis allows us to isolate the effect of direction (increase/decrease) from the magnitude of change after 10Hz stimulation.

We next tested the hypothesis that baseline evoked (CCEP) power in a given time window relates to the strength of pre/post spectral change in that same time window. For example, we predicted that the observation of evoked 10-50ms alpha baseline power in a brain region would yield strong post-stimulation changes in that brain region specifically in alpha power and during the 10-50ms CCEP time window. To test this hypothesis, we compared the t-values in the on-diagonal (i.e. relating pre-stimulation CCEP spectral features to the degree of post-stimulation change, within the same time window and frequency band) and off-diagonal (i.e. relating pre-stimulation CCEP spectral features to the degree of post-stimulation change not in the same time window and/or frequency band) elements using a t-test (ttest2, MATLAB) (FIg 3 Bar plots).

Finally, to study how baseline pre-stimulation time-frequency features influence post-stimulation changes, we performed a 2 (pre, post stimulation CCEP) x 2 (time window, frequency band) ANOVA with follow up post-hoc statistical testing (Tukey’s procedure) (Fig 5, Table 1).

**Fig 5:**
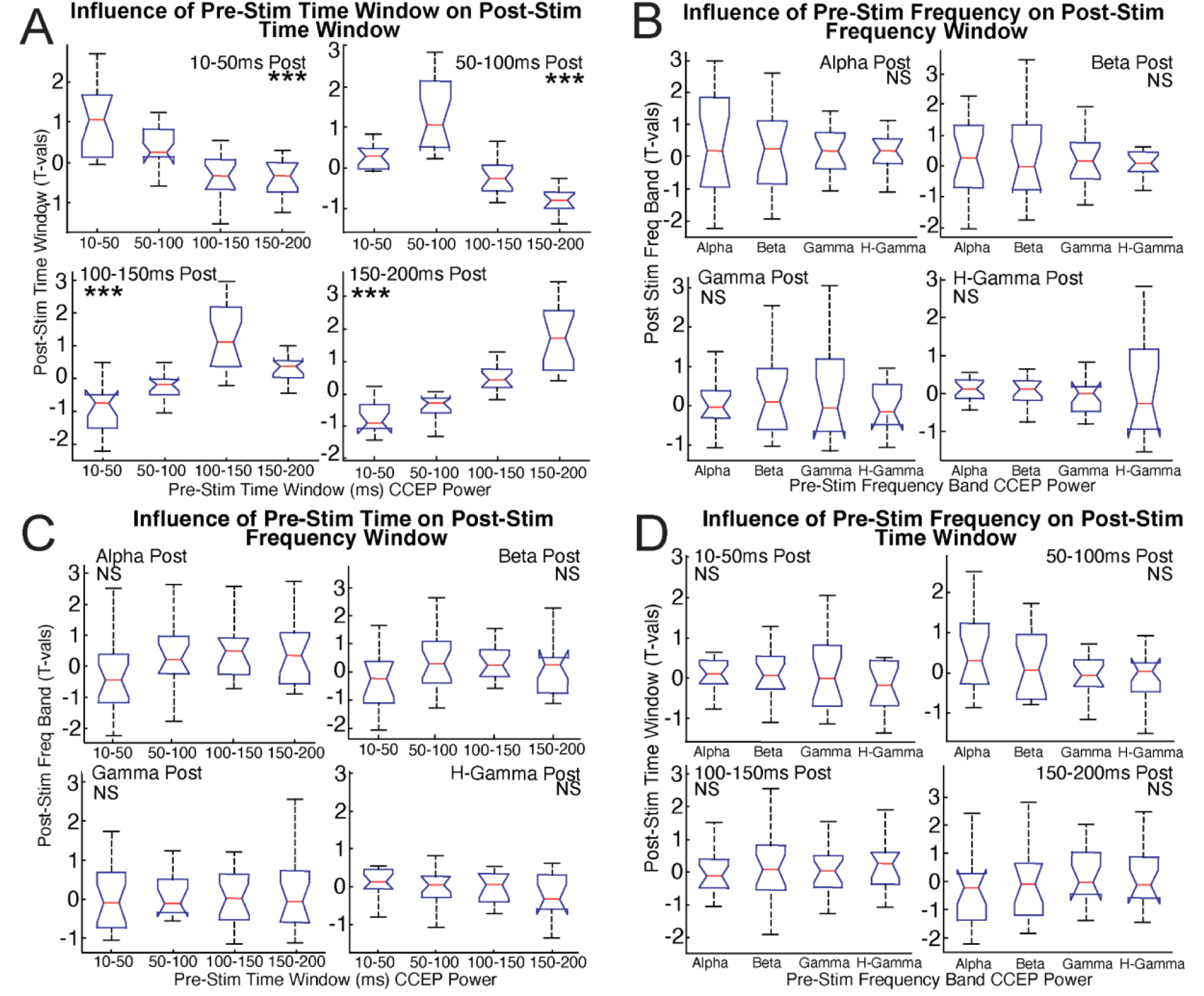
Post-stimulation spectral change is driven by temporally specific baseline CCEP power. A) Group statistical comparison of baseline CCEP power compared to pre/post CCEP power change at each latency. Asterisks represent post-hoc pairwise statistical tests (* = p<0.05 and ** = p<0.01). B) Group statistical comparison of baseline CCEP power compared to pre/post CCEP power change in each frequency. C) Group statistical comparison of baseline CCEP power frequency compared to pre/post CCEP power change in latency . D) Group statistical comparison of baseline CCEP power latency compared to pre/post CCEP power change in each frequency.

**Table 1:** Statistics for Fig 5 - Post-stimulation spectral change is driven by temporally specific baseline CCEP power. Four ANOVA statistics are computed: Mean squared from columns, mean squared from error, the F statistic and the corresponding P value (Prob>F) for the following comparisons: A) Group statistical comparison of baseline CCEP power compared to pre/post CCEP power change at each latency. B) Group statistical comparison of baseline CCEP power compared to pre/post CCEP power change in each frequency. C) Group statistical comparison of baseline CCEP power frequency compared to pre/post CCEP power change in latency . D) Group statistical comparison of baseline CCEP power latency compared to pre/post CCEP power change in each frequency.

|  |  |  |  |  |  |
| --- | --- | --- | --- | --- | --- |
| A | Comparing the effect of post-stim time windows on pre-stim time windows |  |  |  |  |
|  | Post-Stim Time Window | MS from columns | MS from error | F stat | Prob>F |
|  | 10-50 ms | 7.1750 | 0.4316 | 16.63 | 5.63e-08 |
|  | 50-100 ms | 12.8492 | 0.3009 | 42.71 | 6.72e-15 |
|  | 100-150 ms | 12.5440 | 0.5464 | 22.96 | 5.08e-10 |
|  | 150-200 ms | 19.8220 | 0.4452 | 44.53 | 2.87e-15 |
| B | Comparing the effect of post-stim frequency band on pre-stim frequency band |  |  |  |  |
|  | Post-Stim Frequency Band | MS from columns | MS from error | F stat | Prob>F |
|  | Alpha | 0.1296 | 1.4407 | 0.09 | 0.9653 |
|  | Beta | 0.2390 | 1.2563 | 0.19 | 0.9026 |
|  | Gamma | 0.3958 | 0.9295 | 0.43 | 0.7352 |
|  | H-Gamma | 0.1425 | 0.6681 | 0.21 | 0.8868 |
| C | Comparing the effect of post-stim frequency band on pre-stim time windows |  |  |  |  |
|  | Post-Stim Frequency Band | MS from columns | MS from error | F stat | Prob>F |
|  | Alpha | 2.5800 | 1.3181 | 1.96 | 0.1300 |
|  | Beta | 1.8144 | 1.1776 | 1.54 | 0.2123 |
|  | Gamma | 0.0931 | 0.9447 | 0.1 | 0.9605 |
|  | H-Gamma | 0.4112 | 0.6547 | 0.63 | 0.5997 |
| D | Comparing the effect of post-stim time window on pre-stim frequency band |  |  |  |  |
|  | Post-Stim Time Window | MS from columns | MS from error | F stat | Prob>F |
|  | 10-50 ms | 0.7962 | 0.7505 | 1.06 | 0.3725 |
|  | 50-100 ms | 0.3926 | 0.9236 | 0.43 | 0.7357 |
|  | 100-150 ms | 0.2567 | 1.1608 | 0.22 | 0.8813 |
|  | 150-200 ms | 0.4188 | 1.4153 | 0.3 | 0.8282 |

## Results

### Widespread and frequency-specific changes observed after stimulation

First we asked if there were frequency-specific changes in the evoked response after stimulation, and if so, how they differed across various frequencies. Overall, we observed low frequency power (alpha and beta) to be more likely to change after stimulation (Fig 2B,C). Across all time windows, the average group-level percentage of significant channels was 12.87% in the alpha frequency and 7.63% in the beta frequency, while in the gamma and high gamma frequencies, the average was 1.94% and 0.97% respectively(Fig 2B). In contrast, beta and gamma power exhibited the strongest magnitude of change after stimulation. This change was a net suppression of power and most notable in early time windows (Fig 2D, E). In later time windows, a much smaller magnitude of change was observed, across frequency bands. This indicates that the magnitude of the net change across all significant channels was the greatest in this specific time window and frequency band combination. In summary, we observed that low frequency power to be more likely to change after stimulation, while high frequency power in early time windows demonstrated stronger change after stimulation.

### Regions with stronger baseline power are more likely to undergo post-stimulation changes

First, we asked if and how pre-stimulation evoked spectral features relate to post-stimulation evoked spectral changes, predicting that pre-stimulation evoked power correlated with the degree of post-stimulation evoked spectral change. To understand this relationship, we extracted and compared evoked power features in different time windows and frequency bands for pre- and post-stimulation (see Methods and Fig 1). Across patients, we observed that regions with stronger (positive or negative) pre-stimulation power were more likely to undergo a greater post-stimulation spectral change (Fig 2A, 3, 4). Future sections below expand upon this relationship.

### Post-stimulation spectral changes are specific to pre-stimulation evoked power

Next, we explored the degree to which post-stimulation changes are predicted by baseline spectral features. We hypothesized that the specific baseline time-frequency features of the CCEP would strongly relate to post-stimulation effects. In other words, for a given frequency band in a specific time window of the CCEP, baseline spectral power would predict post-stimulation change in the same frequency band in the same time window. On the group level, across all frequencies and time windows, pre-stimulation features were strongly associated with post-stimulation features (note the significant values in the on-diagonal in Fig 3; T_ondiag vs off diag_ (256) = 12.49, p = 3.0448e-28). This effect was consistent in alpha (T_ondiag vs off diag_ (16) = 7.1, p = 1.70e-11), beta (T_ondiag_ = 2.6712, T_offdiag_ =-0.41556, T_ondiag vs off diag_ (16) = 7.24, p = 7.47e-12), gamma (T_ondiag_ = 2.3223, T_offdiag_ = -0.32725, T_ondiag vs off diag_ (16) = 6.81, p = 9.1124e-11), and high gamma (T_ondiag_ = 2.4445, T_offdiag_ = -0.6068, T_ondiag vs off diag_ (16) = 9.25, p = 1.90e-17), as well as in 10-50ms, (T_ondiag_ = 2.1527, T_offdiag_ = 0.67277, T_ondiag vs off diag_ (16) = 3.64, p = 3.3e-4), 50-100ms (T_ondiag_ = 2.5263, T_offdiag_ = 0.90769, T_ondiag vs off diag_ (16) = 3.54, p = 4.9e-4), 100-150ms (T_ondiag_ = 2.575, T_offdiag_ = 0.77392, T_ondiag vs off diag_ (16) = 4.53, p = 1e-5), and 150-200ms (T_ondiag_ = 2.9066, T_offdiag_ = 1.3648, T_ondiag vs off diag_ (16) = 3.92, p = 1.2e-4) time windows. In summary, we observed a strong relationship between baseline power and the degree of post-stimulation changes across frequencies and latencies.

### Direction of baseline evoked power modulates post-stimulation changes in evoked power

Next, we asked if the direction of baseline evoked CCEP power (positive or negative power relative to spontaneous baseline activity) differentially affected post-stimulation changes. We hypothesized that the direction of this baseline evoked power may be an important contributor. To explore this relationship, for each frequency band and time window we stratified brain regions to those with positive and negative baseline evoked power (see Methods and Fig 1 for details). Similar to the non-stratified case (Fig 3), we observed that post-stimulation spectral changes were strongly related to direction of pre-stimulation evoked power (Fig 4B,F B-C; positive baseline features: T_on vs off_ = 12.05; T_on_(16), T_off_ (240); p < 0.001; Fig 4C,G negative baseline features: T_on vs off_ = -4.87; p < 0.001). However, these changes, were specific to the direction of baseline evoked time window and frequency features.

**Fig 4:**
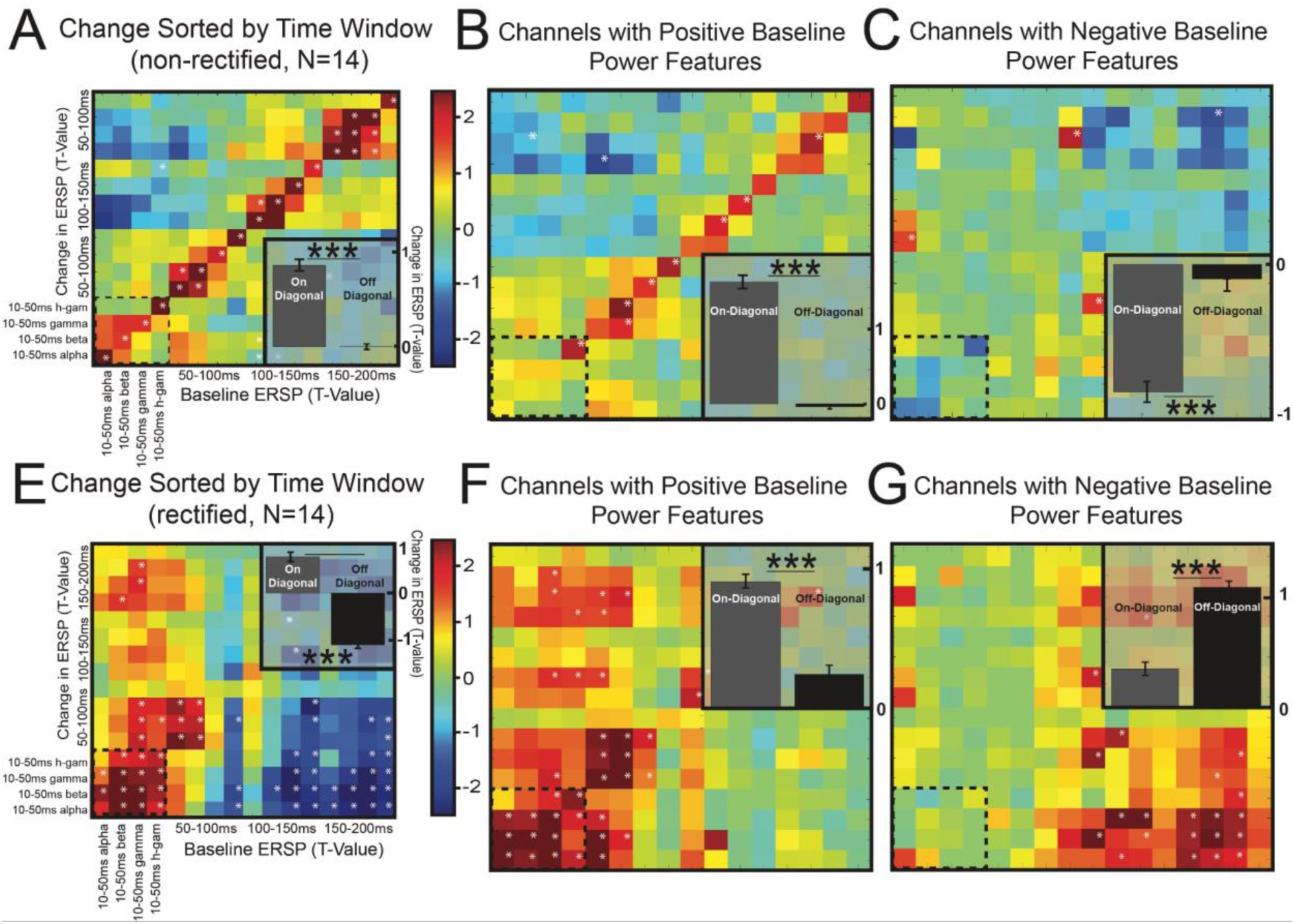
Post-stimulation spectral change is driven by baseline evoked spectral features. Group quantification broken down by the direction of baseline power and sorted by time window. A) Comparison of baseline to pre/post change in ERSP using all channels (same as Fig 3A). B, C) Quantification of baseline power vs change in power, broken down by channels with (B) positive and (C) negative baseline evoked power. E) Relationship between baseline and pre/post change in time-frequency features, but here resulting t-value from pre vs post-stimulation (first t-test) is rectified. F, G (rectified)) Quantification of baseline power vs change in power, broken down by channels with (F) positive and (G) negative baseline evoked power with resulting t-values from pre vs post-stimulation rectified. * in heatmap = p <0.05 from bootstrapped null distribution; *p<.05, **p<.01, ***p<.001 (Student’s T-test). Statistical comparisons between on-diagonal and off-diagonal components for the respective heatmaps are shown as in insert. Note the larger number of positive valued pixels in the rectified heatmap – here warm colors denote strong (positive or negative) pre/post change scores – stratified for those channels with positive (Fig 4B, F) and negative (Fig 4C, G) baseline power. Also note the power decrease from pre-stimulation to post-stimulation; that is, channels with a positive baseline became less positive and those with negative baseline became less negative. This is indicated in the change of pixel color from the non-rectified plot to the rectified plot. Specifically, warm pixels stay warm through rectification in the positive baseline case and the cool pixels turn warm through rectification in the negative baseline case. (Fig 4 E,F,G viewed together N=14).

To explore the relationship between the *direction* of evoked power (increase or decrease) and the *time window* of interest (e.g. 10-50ms after the pulse)with post-stimulation change, we rectified the t-test communicating change in power prior to our statistical test comparing baseline power with pre/post change. This rectification (of change) is important because it helps determine the direction of change (see Methods). First,qualitatively we observed brain regions with positive pre-stimulation evoked power in the early evoked time window (<100ms) to be more likely to undergo changes in post-stimulation power (Fig 4F). We also observed a strong association between negative pre-stimulation evoked power in later time windows (>100 ms) and changes post-stimulation (Fig 4G). This differential effect was found to be statistically significant for both positive (T(128)_<100 vs >100_= 8.43; p = 2.6e-15) and negative (T(128)_<100 vs >100_= -10.29; p = 5.3e-21) pre-stimulation power. Comparing the non-rectified and rectified heatmaps (see Fig 4), one can observe that regions with positive pre-stimulation power increased demonstrated an increase in post-stimulation power (become more positive) in later time windows (>100ms) and a decrease in post-stimulation (become less positive) in early time windows (<100ms). In contrast, regions with negative pre-stimulation power demonstrated an increase in power post-stimulation (become more negative) in early time windows and a decrease in power post-stimulation (become less negative) in later time windows. In summary, we observed that regions exhibiting positive pre-stimulation evoked power in early time windows and negative evoked power in later time windows were both likely to undergo spectral changes after stimulation.

### Post-stimulation change is driven by temporally specific baseline power

Finally, motivated by the by the strong relationship between pre-stimulation baseline power and post-stimulation changes in frequency bands and time windows other than the baseline evoked power in matching frequency band and time window (off-diagonal patterns in Fig 3 F-J), we investigated whether the spectral or temporal nature of the baseline CCEP was more strongly related to spectral changes observed after stimulation. To evaluate this, we stratified the data based on time window and frequency band of the pre- and post-stimulation evoked power and performed an analysis of variance (ANOVA, see Methods for details). Overall, we found that the temporal specificity of baseline power was a more significant driver of post-stimulation changes than baseline frequency (Fig 5) (Average f-stat (Table 1): 31.71 (for the influence of pre stimulation time windows on post stimulation time windows), 0.23 (for the influence of pre stimulation frequency bands on post stimulation frequency bands), 1.06 (for the influence of pre stimulation time windows on post stimulation frequency bands), 0.50 (for the influence of pre stimulation frequency bands on post stimulation time windows)). The effect of stimulation was stronger when examining the same pre-stimulation and pre/post change time window (e.g. 10-50ms baseline compared to 10-50ms pre/post change) compared to when the baseline and change time windows were not aligned (e.g. 10-50ms baseline compared to 50-100ms pre/post change) (Fig 5A, see Table 1 for statistics). In other words, baseline power in a specific time window (eg. 10-50ms) was associated with pre/post change in the same time window (10-50ms) across frequency bands, more strongly than in other time windows. The same was not true when stratifying by frequency bands. That is, no frequency band was associated with pre/post change in any specific frequency band more strongly than others across time windows of the CCEP (Fig 5B, see Table 1 for statistics). Moreover, no frequency band was associated with pre/post change for any specific time window and time window predicted change for any specific frequency band (Fig 5C,D). In summary, we observed that baseline CCEP power in a specific time window was strongly associated with pre/post change in the same time window more strongly than for other time windows. This relationship was not observed in the frequency domain.

### Post-stimulation spectral change patterns are independent of stimulus location

To determine the effect of stimulation location on these results, we stratified results based on stimulation location: premotor (N=6), parietal (N=5), and temporal (N=3) cortex (Table S1). We observed consistent findings across stimulation locations (Fig 5, 6). The prominent effects observed on the diagonal across stimulation sites (Fig 5A-C left panels) were consistent with the notion that post-stimulation changes are specific to pre-stimulation time window and frequency band. Moreover, group maps across all stimulation sites exhibited many positive pixels in the rectified heatmaps stratified by pre-stimulation direction of baseline power, indicating that regions with stronger baseline power had a greater post-stimulation power decrease (Fig 5D-I, right panels). Generally, we also observed the relationship between baseline evoked power direction

and latency and the corresponding post-stimulation change as described above, though this finding was qualitatively more robust for parietal and temporal cortex than premotor (Fig 5). Finally, we also observed baseline power in specific time windows associated with post-stimulation change, in line with our observation in the full group case, that even across brain regions post-stimulation change was driven by temporally specific baseline power more so than by its specific spectral band (Fig 6, Table S1). In summary, findings presented above (Figs 2-4) were observed to be independent of stimulation site.

**Fig 6:**
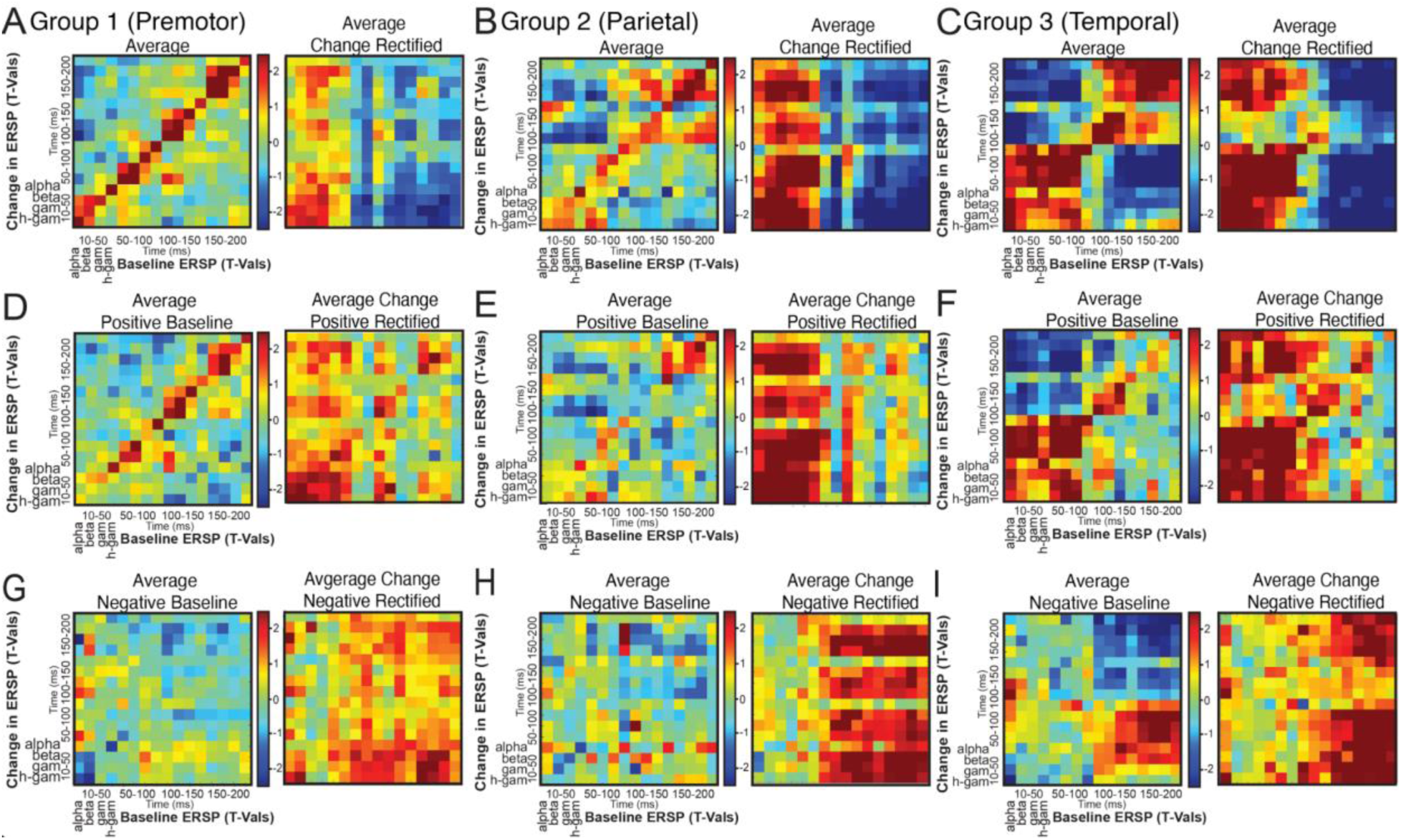
Post-stimulation spectral change patterns are independent of stimulus location. Group quantification for each stimulation location (A, D, G) Premotor, (B, E, H) Parietal and (C, F, I) Temporal. Data are sorted by direction baseline CCEP power (polarity) and time window comparing baseline to pre/post change using (A-C, left) all channels (D-F, left) positive baseline evoked power and (G-I, left) negative baseline evoked power. Also shown is the relationship between baseline and post-stimulation change, but here resulting statistic from pre vs post-stimulation (first t-test) is rectified for (A-C right) all channels with positive baseline evoked power (D-F, right) and negative baseline evoked power (G-I, right).

## Discussion

### Summary of findings

This paper investigates how evoked oscillatory activity changes after 10Hz direct electrical stimulation (Fig 1). Our findings were as follows: Across regions, 10Hz direct electrical stimulation was more likely to modulate evoked alpha power across channels, frequency bands and time windows while stimulation modulated beta/gamma power more strongly than other frequencies (Fig 2). Across patients, regions with stronger evoked baseline power were more likely to undergo greater spectral changes after stimulation (Fig S1A, 2A). These post-stimulation plasticity effects demonstrated specificity in the frequency and time domain (Fig 3) and was driven by an interaction between direction of baseline activity and temporal window (Fig 4). The time window of evoked pre-stimulation power was a stronger predictor of post-stimulation change than frequency (Fig 5, Table 1). The above results were found to be independent of stimulation location (Fig 6, 7, Table S1). Together, these findings demonstrate that time-frequency features before stimulation are strongly associated with oscillatory changes after stimulation.

**Fig 7:**
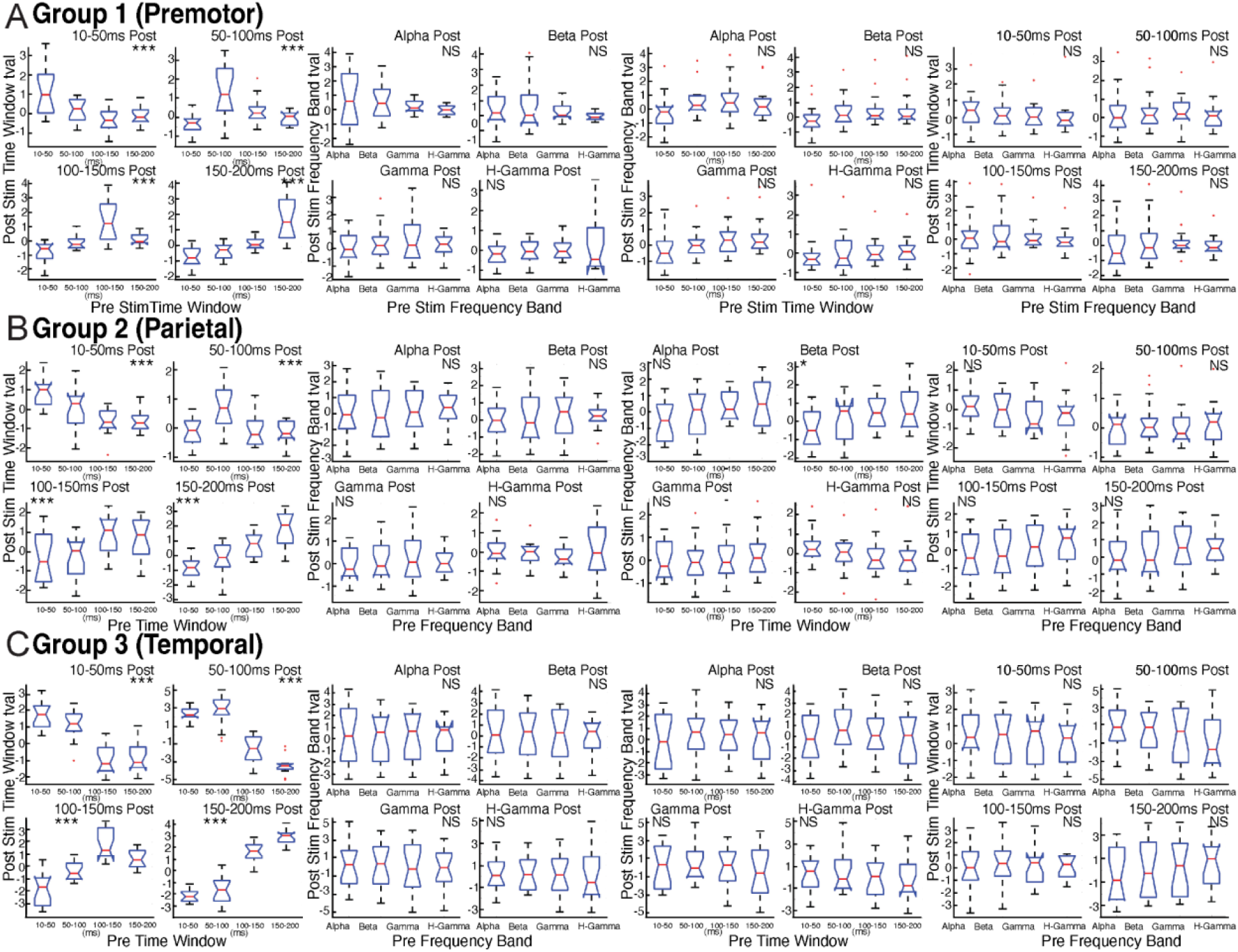
Post-stimulation spectral change is driven by temporally specific baseline CCEP power and is independent of stimulation location. ANOVA statistics for each stimulation location A) Premotor, B) Parietal and C) Temporal. Left: Group statistical comparison of baseline power compared to post-stimulation power change at each latency. Second to left: Asterisks represent post-hoc pairwise statistical tests (* = p<0.05 and ** = p<0.01). Group statistical comparison of baseline power compared to post-stimulation power change in each frequency (second from left). Third to left: Group statistical comparison of baseline power frequency compared to post-stimulation power change in latency. Right: Group statistical comparison of baseline power latency compared to post-stimulation power change in each frequency.

In our previous studies, we showed that post-stimulation CCEP changes in the time domain could be predicted by baseline anatomical and functional (CCEP) connectivity profiles (Keller 2018). We next demonstrated additionally that neural activity *during* simulation was an indicator of post-stimulation changes (Huang 2019). Here we extend our analysis to the frequency domain, a physiological dimension that appears critical for the brain’s functioning (Buzsáki & Draguhn, 2004). We observed that evoked baseline power in a given time window and frequency band were most strongly associated with short-term plasticity effects in the same time window and frequency band (Fig 2). While only a few studies have examined neural changes in the time-frequency domain after repetitive stimulation (Woźniak-Kwaśniewska et al., 2014, Brignani et al., 2008, Fox et al., 2020 & Veniero et al., 2011), our results are broadly in line with other studies that demonstrate a frequency dependent change in neural activity and connectivity after stimulation (Fox et al., 2020).

We also extend our work to demonstrate that the direction of baseline power (positive or negative) is strongly associated with post-stimulation effects. We observed regions with positive baseline power in earlier CCEP time windows and negative baseline power in later CCEP time windows were more likely to increase in magnitude after repetitive stimulation (Fig 3). However, due to the myriad neural mechanisms that can produce the same direction of change in power, evoked with single pulses, both at baseline and after repetitive stimulation, results should be interpreted with caution. While intriguing, interpreting post-stimulation power changes in brain regions and frequencies that exhibit negative evoked power at baseline is difficult at best, and highlights the need for a deeper understanding of the neural mechanism underlying these changes.

Moreover Hebbian theory suggests that repeatedly activating regions that are functionally connected (‘fire together’) cause eventual neural plasticity (‘wire together’) (Hebb, 1949). Put another way, Hebbian theory describes a basic mechanism for synaptic plasticity, in which repeated stimulation of postsynaptic cells by their presynaptic counterparts results in synaptic changes. This phenomenon is well described in animals where electrical stimulation applied at specific frequencies have been shown to strengthen excitatory synapses in mice (Vlachos et al., 2012, Bliss & Lomo, 1973, Bear & Abraham, 1996, Kirkwood et al., 1996 & Malenka & Bear, 2004, Kronberg et al., 2020). However, critically this effect remains poorly understood in humans. Findings presented here demonstrating that regions with stronger baseline power (analogous to ‘wire together’) were more likely to be correlated with greater post-stimulation change (analogous to ‘fire together’). These findings exhibited a time window / frequency band specificity of post-stimulation effects. Together, this work provides initial support for the observation of a Hebbian-like learning to occur in humans. Furthermore, in contrast to animal studies which focus in the time series domain, this work explores similar phenomena in the frequency domain. As such, these findings help improve our understanding of human brain plasticity and in the future may lead to more effective brain stimulation techniques.

Finally these results introduce ideas that are clinically translatable in conjunction with the work in our previous studies. In (Keller et al., 2018) we showed that neural plasticity (i.e. excitability changes) were maintained after stimulation for up to 15 minutes. The (Huang et al., 2019) paper built upon this investigating intra-stimulation brain changes with findings that could potentially be used for real-time implementation. In this study we confirm the need to focus on the time window and frequency band of the evoked potential by demonstrating that post-stimulation plasticity was driven by specific time-frequency baseline features and also that the temporal specificity of baseline power was more indicative of plasticity than the spectral specificity.

### Future directions

The overarching goal of this work was to investigate the neural mechanisms underlying a Hebbian-like plasticity induced by 10Hz direct electrical stimulation. The longer-term goal is to translate these ideas noninvasively to simultaneous TMS and scale EEG (TMS-EEG) in an effort to predict which regions and frequency content will change. By doing so, one could modify (and therefore personalize) the treatment target or stimulation pattern to optimize predicted change prior to treatment. As such testing the robustness of these findings on subsequent electrophysiological datasets from noninvasive and invasive stimulation and recording modalities will improve our understanding and study the generalizability of these findings, increasing translation potential. Moreover, by examining the spatiotemporal neural effects that occur *during* stimulation (intra-stimulation effects) we can in a more granular manner test if these Hebbian-style neural effects generalize to the time periods during and after stimulation. Hence, several steps are needed for this important translation: (1) A better understanding of the temporal evolution of these spectral changes, both *during* and *after* repetitive stimulation; (2) A better understanding of the intracranial acute neural effects and dynamics of change after stimulating at different frequencies (i.e. 1Hz vs 5Hz vs 10Hz) and patterns (i.e. theta burst stimulation); (3) Examining the time-frequency response during and after applying stimulation patterns thought to be ‘excitatory’ (e.g. 10Hz, 100Hz, theta burst) and ‘inhibitory’ (e.g. 1Hz), and testing the consistency of these neural effects across stimulation location (4); Performing further statistical analyses by clustering neural effects in space and time; that is, finding those subnetworks that are co-modulated similarly during and after repetitive stimulation. Future work will need to translate these findings to noninvasive neuromodulatory treatments such as TMS (Jeffrey B. Wang et al., 2022) in order to move towards clinical translation.

### Limitations

The generalizability of this study is limited by various considerations. Most importantly, electrical stimulation and recordings were performed on epilepsy patients. As a result, findings are limited by patient etiology and electrode placement. As indicated in previous work, local and global brain connectivity patterns can be disrupted in this patient population (Pereira et al., 2010, Bettus et al., 2011 & Pittau et al., 2012). Furthermore, it was difficult to perform a more thorough assessment of stimulation parameters (site, frequency, intensity) due to experimental time constraints (allotted < 1 hr per patient). Secondly, single electrical pulses used to derive CCEPs resulted in prominent harmonics directly after stimulation, resulting in some ambiguity in the frequency that changed after stimulation (i.e. spectral change at 80Hz could be due to harmonics of a 40Hz change or a primary change at 80Hz)). Finally, this study focused on focal electrical stimulation in an attempt to learn about the underlying neural mechanisms of noninvasive TMS. We applied direct electrical stimulation in this study because the goal was to provide a ‘ground truth’ understanding of the neural effects of repetitive stimulation, and electrical stimulation is focal and without perceptual effects. But of course electrical stimulation is not TMS and both likely activate brain tissue differently (Borchers et al., 2011).

## Conclusions

In this paper, we investigated the effect of electrical patterned stimulation on neural oscillations in the human brain. This coupling of baseline oscillatory activity with post-stimulation changes in intracranial data provides important insight into plasticity induction in humans. This work opens new lines of scientific investigation including personalizing pre-stimulation neural features to maximize post-stimulation changes for neuropsychiatric disorders.

## Acknowledgments

The authors are grateful for the participants that dedicated their time to this research. All authors discussed the data, analysis, and methods and contributed to the manuscript. The authors are enormously indebted to the patients that participated in this study, as well as the nursing and physician staff at North Shore University Hospital (Manhassat, NY) and the National Institute of Clinical Neurosciences (Budapest, Hungary) as well as to Boglárka Hajnal, László Entz and Dániel Fabó (National Institute of Clinical Neuroscience, Budapest, Hungary) and Jose L. Herrero, Stephan Bickel, and Ashesh Mehta (Neurosurgery, Hofstra Northwell School of Medicine and Feinstein Institute for Medical Research) for their contribution to the study design and data collection. We thank Gayathri Ganesan (Stanford University) for help with preliminary data analysis. CJK is funded by R01MH129018, R01MH126639, R01MH132074, and the Burroughs Wellcome Fund Career Award for Medical Scientists.

## Author contributions

C.J.K. designed and performed experiment; Y.H. preprocessed the data. S.M. and N.K. analyzed the data and created figures; S.M. and C.J.K. wrote the paper. All authors reviewed and contributed to the manuscript.

## Supplementary Figures and Tables

**Table S1:** Statistics for Figure 6. Post-stimulation spectral change is driven by temporally specific baseline CCEP power independent of stimulation location. The F statistic and the corresponding P value (Prob>F) are calculated for each stimulation location and time window-frequency band pair.

| Group 1 - RPM |  |  | Group 1 - Parietal |  |  | Group 3 - Temporal |  |  |
| --- | --- | --- | --- | --- | --- | --- | --- | --- |
| Post Time windows for pre | time windows |  | Post Time windows for pre | time window |  | Post Time windows for pre | time windows |  |
| Post Time window | F stat | Prob>F | Post Time window | F stat | Prob>F | Post Time window | F stat | Prob>F |
| 10-50 ms | 11.02 | 7.31E-06 | 10-50 ms | 14.22 | 4.15E-07 | 10-50 ms | 36.99 | 1.15E-13 |
| 50-100 ms | 14.3 | 3.87E-07 | 50-100 ms | 9.25 | 4.05E-05 | 50-100 ms | 87.99 | 6.06E-22 |
| 100-150 ms | 17.24 | 3.46E-08 | 100-150 ms | 5.65 | 1.8 e-3 | 100-150 ms | 35.25 | 2.91E-13 |
| 150-200 ms | 26.88 | 3.82E-11 | 150-200 ms | 20.04 | 4.07E-09 | 150-200 ms | 131.22 | 2.57E-26 |
| Post frequency band for | pre frequency | band | Post frequency band for | pre frequency | band | Post frequency band for | Pre frequency | band |
| Post Frequency Band | F stat | Prob>F | Post Frequency Band | F stat | Prob>F | Post Frequency Band | F stat | Prob>F |
| Alpha | 1.14 | 0.341 | Alpha | 0.21 | 0.8875 | Alpha | 0.01 | 0.9976 |
| Beta | 1.09 | 0.3595 | Beta | 0.24 | 0.8671 | Beta | 0.06 | 0.9015 |
| Gamma | 0.69 | 0.5627 | Gamma | 0.29 | 0.8343 | Gamma | 0.17 | 0.9169 |
| H-Gamma | 0.66 | 0.5787 | H-Gamma | 0.38 | 0.7694 | H-Gamma | 0.04 | 0.99 |
| Post Frequency band for | pre time | window | Post frequency band for | pre time windows |  | Post Frequency band for | Pre time | window |
| Post Frequency Band | F stat | Prob>F | Post Frequency Band | F stat | Prob>F | Post Frequency Band | F stat | Prob>F |
| Alpha | 1.47 | 0.2321 | Alpha | 2.37 | 0.0791 | Alpha | 0.29 | 0.8289 |
| Beta | 1.37 | 0.2611 | Beta | 3.26 | 0.0277 | Beta | 0.41 | 0.7457 |
| Gamma | 1.75 | 0.166 | Gamma | 0.33 | 0.8013 | Gamma | 0.36 | 0.7837 |
| H-Gamma | 0.22 | 0.8827 | H-Gamma | 1.39 | 0.255 | H-Gamma | 0.34 | 0.797 |
| Post time window for pre | frequency | band | Post time window for pre | Frequency | band | Post time window for pre | frequency | band |
| Post Time window | F stat | Prob>F | Post Time window | F stat | Prob>F | Post Time window | F stat | Prob>F |
| 10-50 ms | 0.5 | 0.6823 | 10-50 ms | 0.94 | 0.4269 | 10-50 ms | 0.36 | 0.7792 |
| 50-100 ms | 0.13 | 0.9423 | 50-100 ms | 0.16 | 0.9244 | 50-100 ms | 1.07 | 0.3683 |
| 100-150 ms | 0.36 | 0.7841 | 100-150 ms | 1.2 | 0.3166 | 100-150 ms | 0.09 | 0.963 |
| 150-200 ms | 0.53 | 0.6632 | 150-200 ms | 0.5 | 0.6844 | 150-200 ms | 0.47 | 0.7064 |

**Figure S1:**
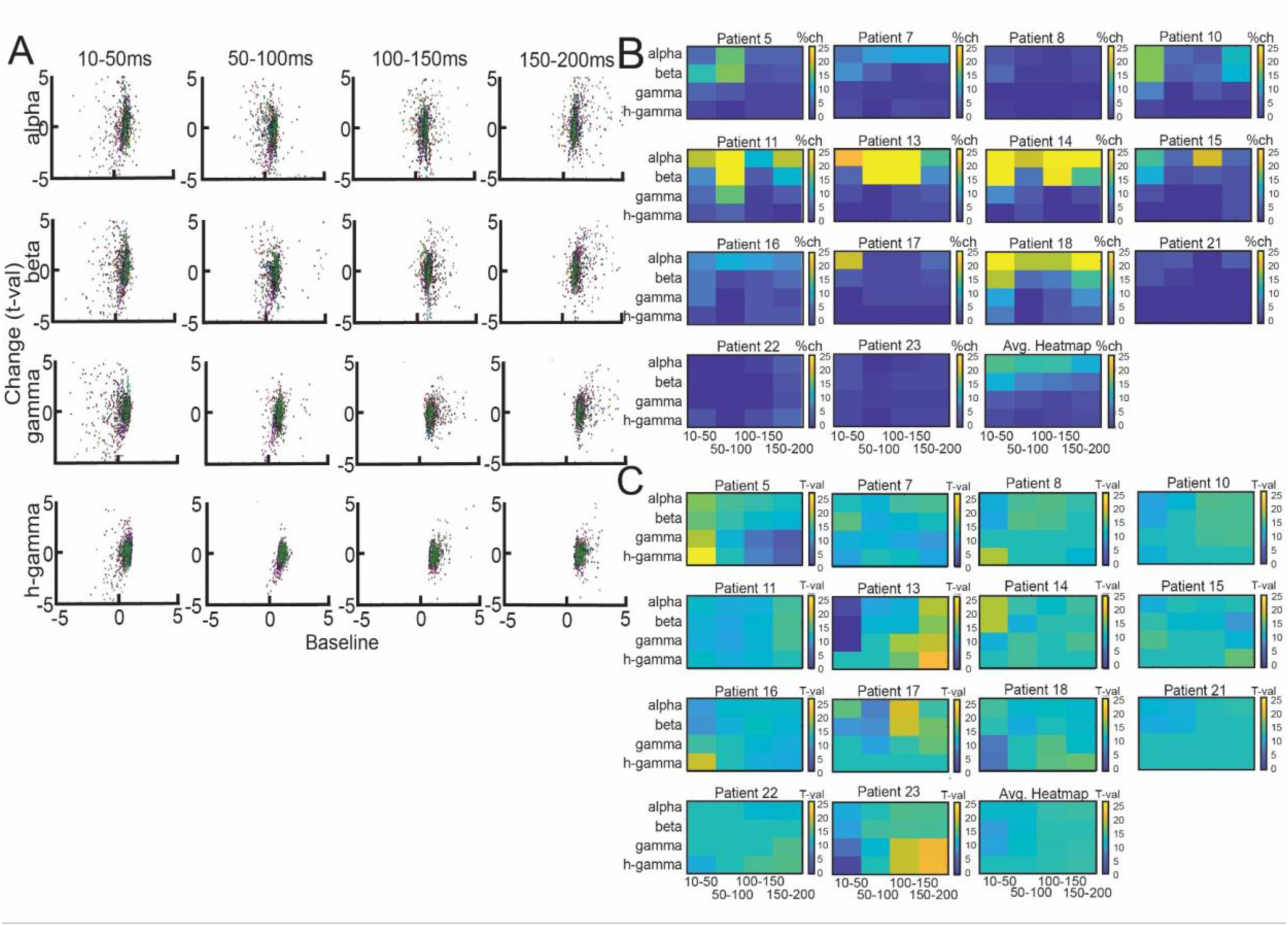
Relationship between spectral features, sensitivity to change and magnitude of change in neural oscillatory activity after repetitive stimulation for all patients. A) Distribution of channels for all patients per frequency band and time window depicting relationship between baseline power and change in power. 2 B, C) Heatmap and bar plots showing sensitivity to change (characterized by number of significant channels (p<0.05) for each stratified frequency band and time window across all subjects. 2 D, E) Heatmap and bar plots showing net magnitude of change across all significant channels across all subjects.

**Figure S2:**
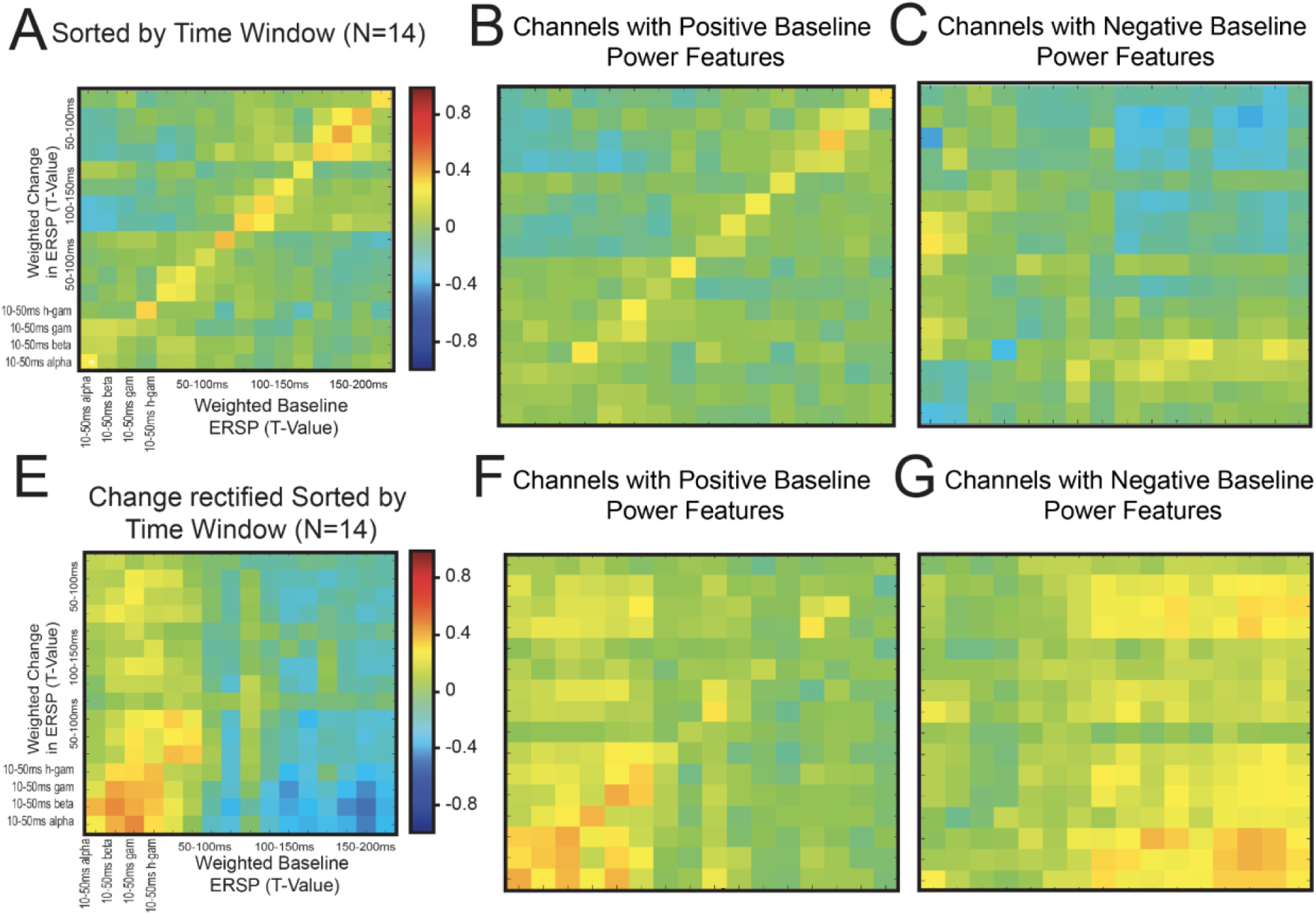
Weighted average change in neural excitability is driven by baseline CCEP spectral and temporal features. Group quantification broken down by direction of baseline CCEP power and sorted by time window, with a weighted average across all patients. A) Comparison of baseline to pre/post change in ERSP using all channels. B, C) Map of baseline power vs weighted average (across patients) change in power, broken down by channels with (B) positive and (C) negative baseline evoked power. E) Relationship between baseline and pre/post change in time-frequency features, but here resulting weighted average t-value from pre vs post-stimulation (first t-test) is rectified. F, G) Map of baseline power vs change in power, broken down by channels with (F) positive and (G) negative baseline evoked power with resulting weighted average t-values from pre vs post-stimulation rectified. * in heatmap = p <0.05 from bootstrapped null distribution; *p<.05, **p<.01, ***p<.001 (Student’s T-test). Statistical comparisons between on-diagonal and off-diagonal components for the respective heatmaps are shown as in insert. (Analogous to Fig 4 in main text)

**Figure S3:**
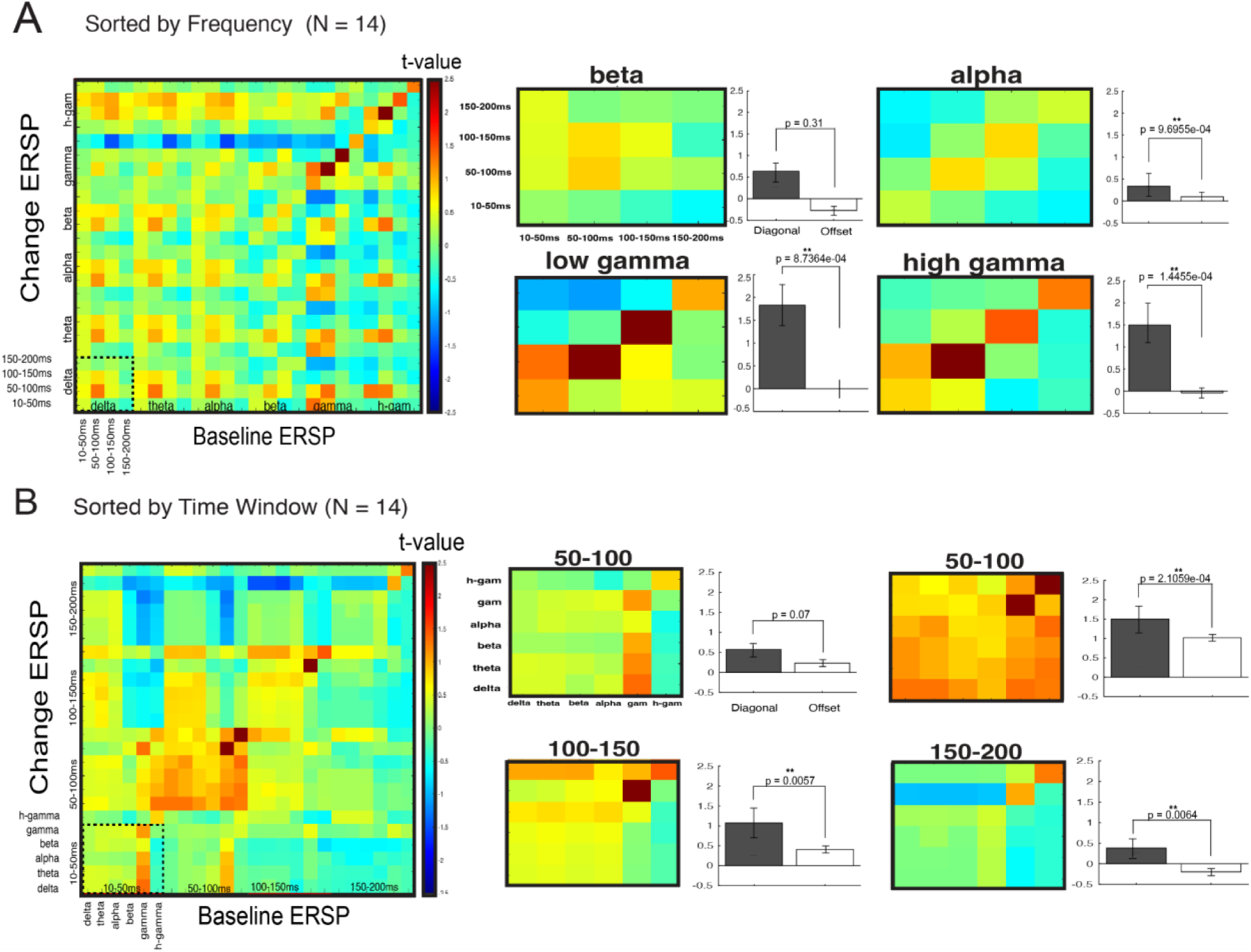
Baseline spectral features predict change in neural excitability-after repetitive stimulation using the spectrogram approach. A) Group quantification relating baseline ERSP and change in ERSP. Axes are sorted by (A) time window and (F) frequency. Note the strong effect between baseline spectral-temporal features and the magnitude of change of those features after 10Hz stimulation, as quantified by the diagonal. *p<0.05 from bootstrapped null distribution (see Methods). B-E) Same data from A but visualized within each latency. Bar graphs represent comparison of on-diagonal vs off-diagonal; that is, how the same baseline temporal-spectral feature predicts post-stimulation change. F) Same as A but sorted by frequency band then latency. G-J) Same as B-E but sorted by frequency band. For bar graphs, *p<.05, **p<.01, ***p<.001 (Student’s T-test)

**Figure S4:**
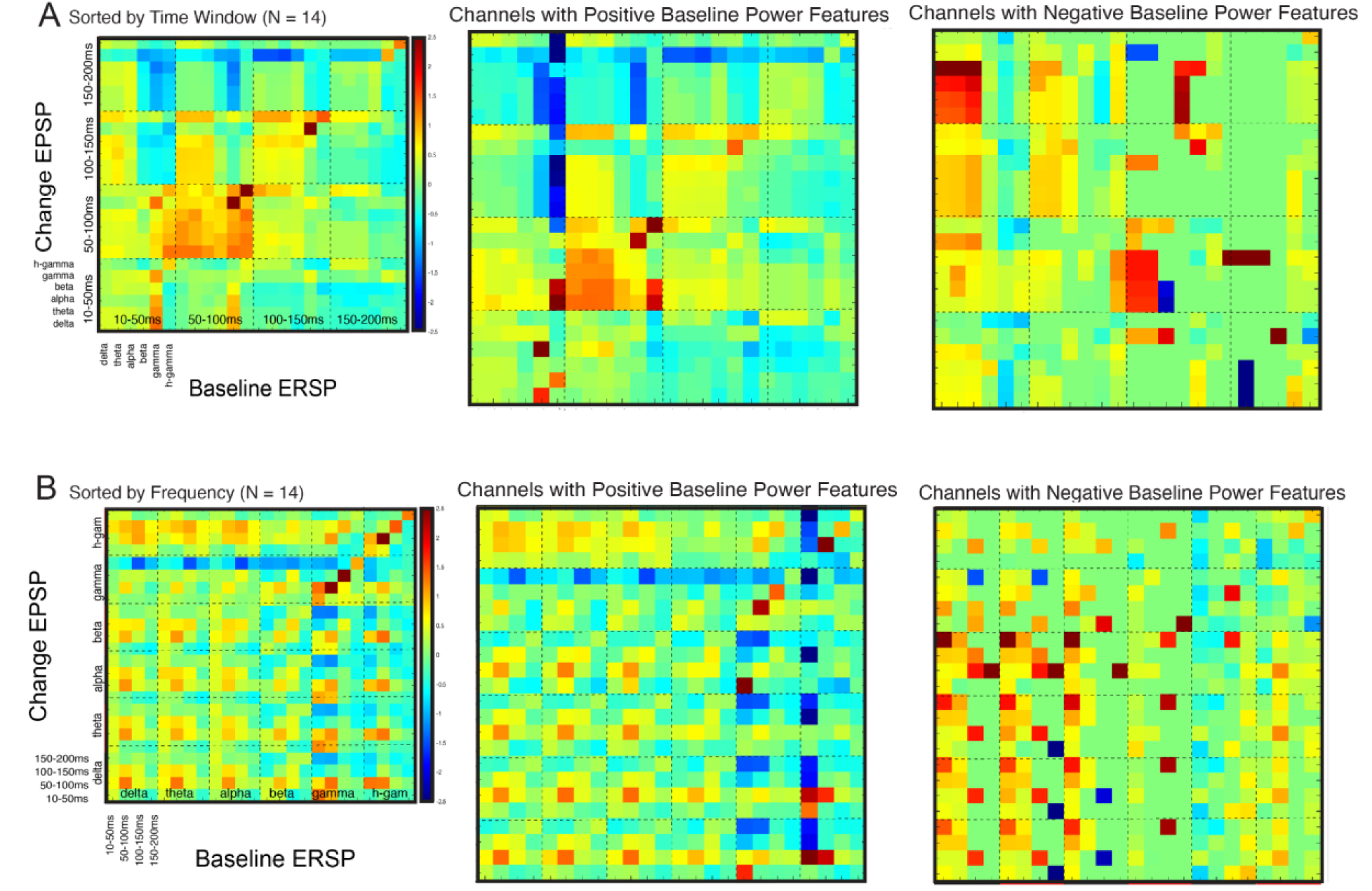
Change in neural excitability is driven by baseline CCEP spectral and temporal features through the spectrogram approach. A) Group quantification broken down by baseline CCEP power polarity (direction of baseline activity) and sorted by time window. *Left:* Comparison of baseline to pre/post change in ERSP using all channels (same as Fig 2A). *Middle/Right:* Map of baseline power vs change in power, broken down by channels with (middle) positive and (right) negative baseline evoked power. B) Relationship between baseline and pre/post change in time-frequency features, but here resulting t-value from pre vs post-stimulation (first t-test) is rectified. * in heatmap = p <0.05 from bootstrapped null distribution; *p<.05, **p<.01, ***p<.001 (Student’s T-test). Statistical comparisons between on-diagonal and off-diagonal components for the respective heatmaps are shown as in insert.

